# Structural Insights Into the Initiation and Elongation of Ubiquitination by Ubr1

**DOI:** 10.1101/2021.04.12.439291

**Authors:** Man Pan, Qingyun Zheng, Tian Wang, Lujun Liang, Junxiong Mao, Chong Zuo, Ruichao Ding, Huasong Ai, Yuan Xie, Dong Si, Yuanyuan Yu, Lei Liu, Minglei Zhao

## Abstract

The N-end rule pathway was one of the first ubiquitin (Ub)-dependent degradation pathways to be identified. Ubr1, a single-chain E3 ligase, targets proteins bearing a destabilizing residue at the N-terminus (N-degron) for rapid K48-linked ubiquitination and proteasome-dependent degradation. How Ubr1 catalyses the initiation of ubiquitination on the substrate and elongation of the Ub chain in a linkage-specific manner through a single E2 ubiquitin-conjugating enzyme (Ubc2) remains unknown. Here, we report the cryo-electron microscopy structures of two complexes representing the initiation and elongation intermediates of Ubr1 captured using chemical approaches. In these two structures, Ubr1 adopts different conformations to facilitate the transfer of Ub from Ubc2 to either an N-degron peptide or a monoubiquitinated degron. These structures not only reveal the architecture of the Ubr1 complex but also provide mechanistic insights into the initiation and elongation steps of ubiquitination catalysed by Ubr1.

## INTRODUCTION

Ubiquitination is involved in a wide range of cellular processes, such as protein homeostasis, cell cycle regulation, transcriptional regulation, and the stress response ^1,2^. In particular, the N-end rule pathway was the first specific pathway of the ubiquitin (Ub) system to be identified ^3^. This pathway determines the rate of protein degradation through recognition of the N-terminal residues termed N-degrons. In eukaryotes, N-degrons are recognized by specific ubiquitin ligases (E3) followed by rapid polyubiquitination of a nearby lysine residue, which marks the protein for degradation by the 26S proteasome ^4–7^. It has been estimated that more than 80% of human proteins can be regulated by the N-end rule pathway ^8^. Misregulation of the N-end rule pathway leads to the accumulation of unwanted proteins and proteotoxicity, which underlies ageing and specific diseases, including neurodegeneration ^9^.

Three branches of the N-end rule pathway exist in eukaryotes, targeting N-terminal arginine residues (Arg/N), proline residues (Pro/N), and acetyl groups (Ac/N) ^10^. In *Saccharomyces cerevisiae* (Baker’s yeast), a single E3 ligase, Ubr1, is responsible for the Arg/N-end pathway, which recognizes two types of N-degrons: one type starts with basic residues (R, K, H), and the other starts with bulky hydrophobic residues (L, F, W, Y, I) ^11^. By contrast, in mammals, multiple Ubr1 homologues participate in the Arg/N-end pathway ^12^. Human Ubr1 shares 20% sequence identity with yeast Ubr1 and is involved in the processes of neurite outgrowth and axonal regeneration ^13^. Mutations to human Ubr1 are associated with the congenital disorder known as Johanson-Blizzard syndrome ^9^. Ubr1 is a single-subunit RING-type E3 ligase with a mass of over 200 kDa ^14^. In vitro, yeast Ubr1 catalyses rapid K48-linked polyubiquitination on substrate proteins that satisfy the N-end rule through the single E2 ubiquitin-conjugating enzyme Ubc2 (also known as Rad6) ^11^, which has been used to prepare ubiquitinated substrates for in vitro assays ^15^. Despite being discovered over 30 years ago ^14^, the overall architecture of Ubr1 remains unknown. More importantly, how Ubr1 installs the first Ub onto the substrate and the subsequent Ub molecules in an efficient and linkage-specific manner remains to be elucidated. Due to the transient nature of the reaction intermediates, visualization of this process is challenging.

Here, we developed chemical strategies to mimic the reaction intermediates of the first and second Ub transfer steps catalysed by yeast Ubr1 and determined two cryo-electron microscopy (cryo-EM) structures of Ubr1 in complex with Ubc2, Ub and either an N-degron peptide or a ubiquitinated N-degron peptide, representing the initiation and elongation steps of ubiquitination, respectively. The overall structures of these two complexes showed remarkable resemblances to that of anaphase-promoting complex/cyclosome (APC/C), a highly conserved multi-subunit E3 ligase involved in eukaryotic cell cycle regulation. Key structural elements, including a Ubc2 binding region (U2BR) and an acceptor Ub binding loop on Ubr1, were identified and characterized, which provided molecular insights into the initiation and elongation steps of ubiquitination catalysed by Ubr1. Ubiquitination underlies many fundamental cellular processes and is also critical for the development of novel therapeutic strategies, such as proteolysis-targeting chimera (PROTAC) ^16,17^. Our chemical approaches provide a general strategy for the structural characterizations of other systems.

## RESULTS

Previous work has shown that an Arg/N-degron peptide consisting of 43 amino acids can be polyubiquitinated at a Lys residue with K48 linkages ^15^. We first synthesized the degron peptide (hereafter referred to as Degron) and a monoubiquitinated Degron (hereafter referred to as Ub- Degron) using solid phase peptide synthesis (**Fig. S1a & b**). For Ub-Degron, the native isopeptide bond was introduced by using a side chain-modified Lys residue through solid phase peptide synthesis. Subsequently, hydrazide-based native chemical ligation (NCL) was conducted to place Ub onto the Lys side chain ^18^ (**Fig. S1b**). To monitor the polyubiquitination reaction, we further labelled both Degron and Ub-Degron with fluorescein-5-maleimide (**Fig. S1c & d**) and confirmed their reactivity in a K48 linkage-specific manner with Ubr1 and Ubc2 in vitro (**Fig. S1e**). Next, we performed single-turnover ubiquitination reactions using Ubc2 loaded with mutant Ub (K48R) at a saturating concentration. The estimated *K*_m_ and *K*_cat_ for Degron were 1.24 ± 0.69 μM and 0.27 ± 0.07 min^-1^, respectively, which represents the initiation step catalysed by Ubr1 (**Fig. S1f**). The single-turnover ubiquitination of Ub-Degron, representing the first elongation step catalysed by Ubr1, showed slightly slower kinetics with an estimated *K*_m_ of 1.63 ± 0.69 μM and *K*_cat_ of 0.17 ± 0.02 min^-1^ (**Fig. S1g**). Notably, this behaviour was different from that of multi-subunit cullin-RING ubiquitin ligase (CRL)-mediated ubiquitination, which is a two-step process involving a slow initiation step followed by rapid K48-specific chain elongation^19,20^.

To capture the initiation step of ubiquitination catalysed by Ubr1, we synthesized a stable complex of Ubc2, Ub and Degron that mimicked the reaction intermediate (**Fig. 1a and Fig. S2a**). A similar design was used in recent studies of the SCF-E3 complex; however, in this work, the Ub moiety was conjugated to the natural ε-amino group of K17 in Degron instead of being directly fused to the N-terminus of the substrate ^21,22^. This stable intermediate mimic was then mixed with Ubr1 in a 1:1.5 molar ratio and incubated on ice for 30 minutes followed by vitrification and single-particle cryo-EM analysis (**Fig. S2c**). A dataset of 10,592 movie stacks was collected and processed following the established workflow in RELION ^23^ (**Fig. S3a**). The final reconstructed map had an overall resolution of 3.3 Å, allowing for de novo model building (**Figs. S3b, 3c, & 4a; Table S1**). Ubr1 is a single-subunit E3 ligase with more than 1,900 amino acids (**Fig. 1b**). Only the Ubr-Box1 structure was determined previously ^24^. To overcome the difficulty during model building, we used the artificial intelligence (AI)-based do novo modelling tool DeepTracer ^25^ to build a starting model of the entire complex (see Methods for details) and manually adjusted and refined the model using COOT ^26^. The final model was refined in real space using PHENIX ^27^ (**Table S2**).

**Figure 1:**
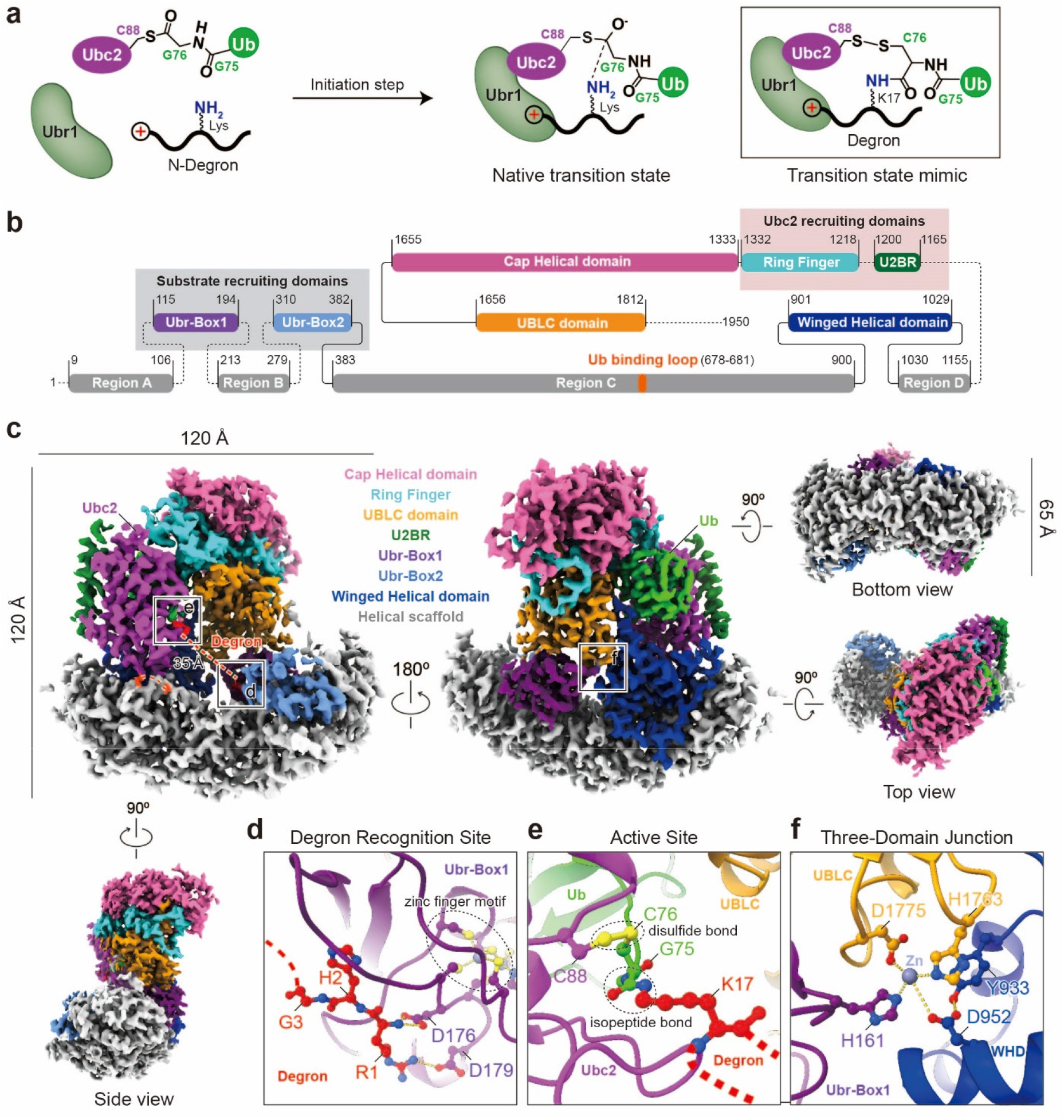
Structure of the initiation complex. **a**, A schematic representation of the transition state of the initiation step. The side chain of a Lys residue on Degron attacks the thioester bond of Ubc2∼Ub. The inset shows the designed intermediate structure mimicking the transition state of the initiation step. **b**, A schematic domain diagram of Ubr1, with residue boundaries indicated. Dotted lines represent unresolved linkers and regions. **c**, Cryo-EM maps of the initiation complex (sharpened using a B factor of −96.5 Å^2^, contour level: 0.03). The colour code of Ubr1 is the same as that in panel **b**. **d,** Molecular interactions between Degron and Ubr-Box1. Hydrogen bonds and electrostatic interactions are represented by dotted lines. **e,** Molecular structure of the active site. **f,** Molecular interactions between UBLC, Ubr-Box1 and the winged helical domain (WHD). Metal coordination bonds and hydrogen bonds are represented by dotted lines. Mutations of the four residues at the interface (H161, Y933, H1763 and D1775) impaired the activity of Ubr1.

The overall structure of the initiation complex, named Ubr1-Ubc2-Ub-Degron, adopted a sailboat-like shape, bearing high resemblance to APC/C ^28,29^, although Ubr1-Ubc2-Ub-Degron was much smaller at 120 Å × 120 Å × 65 Å (**Fig. 1c**). The hull of the “boat” is a large helical scaffold (white-grey) interspaced by three domains: Ubr-Box1 (dark purple), Ubr-Box2 (light blue), and a winged helical domain (dark blue). The helical scaffold is reminiscent of similar bundle repeats found in other E3 ligases, especially the cullin proteins in CRLs ^21,22,30^. We further defined four regions of the helical scaffold based on the interspaced domains (**Fig. 1b, Fig. S5a & b**). Only the structure of Ubr-Box1 has been previously reported ^24^. The other domains were identified using SWISS-MODEL ^31^. The front of the “boat” is where the Ub-loaded Ubc2 is recruited. Ubc2 is primarily bound by a single helix of Ubr1, termed the Ubc2 binding region (U2BR, dark green, **Fig. 1b & c**). A RING finger domain (cyan) follows U2BR and interacts with Ubc2 and the loaded Ub (**Fig. S5d**). A new helical domain termed the cap helical domain (pink) follows the RING finger domain (**Fig. S5c**). The cap helical domain adopted a new fold that did not give similar hits from the DALI server for protein structure comparison ^32^. Finally, the UBLC domain (UBR/Leu/Cys domain) ^33^ (orange) acts like the mast of the “boat” by interacting with Ubr-Box1 and the winged helical domain through a zinc-binding site (**Fig. 1c & f**). Quadruple mutations of the residues involved in this interface (H161A, Y933A, D1175A, and H1763A) greatly impaired the activity of Ubr1 (**Fig. 1f, Fig. S5f**).

The first three residues (RHG) of the Degron peptide were resolved bound to the designated pocket of Ubr-Box1 (**Fig. 1d, Figs. S4a & 5e**) ^24^. The active site of Ub transfer is ∼35 Å away from the C-terminus of Gly3, where Lys17 forms an isopeptide bond with the C-terminal Cys76 of Ub (**Fig. 1e**). The thirteen residues between Lys17 and Gly3 were not resolved, but the distance between them indicated an extended conformation (2.7 Å/residue). Meanwhile, a disulfide bond was formed between the cysteine and the active site of Ubc2 (Cys88), as designed (**Fig. 1e**). Ub is on the backside of the complex and interacts with one of the zinc-binding sites in the RING finger domain through the Ile36 patch (**Fig. S5d**). In addition, the U2BR of Ubr1 forms an extensive interface with the backside of Ubc2 (**Fig. 1c**), reminiscent of the Ube2g2 binding region (G2BR) in Gp78, an E3 ligase involved in endoplasmic reticulum-associated degradation (ERAD) ^34^, and the Ubc7 binding region (U7BR) in Cue1p, a component of several E3 complexes involved in ERAD ^35^. Together, the noncovalent interactions between Ubr1, Ubc2, and Ub position the Ubc2∼Ub thioester bond for nucleophilic attack by Lys17 on Degron. In summary, this complex structure shows how Ubr1 recruits Ubc2∼Ub and facilitates the transfer of Ub to a specific Lys residue on the Degron peptide; that is, the initiation step.

Once the first Ub is installed on the substrate, the subsequent elongation of the Ub chain cannot occur without rearrangement of the structure. To understand the process and capture the intermediate state of the elongation step, we designed another stable complex mimicking the transition state (**Fig. 2a**). The C-terminus of donor Ub was linked to both Cys88 of Ubc2 (active site) and C48 of acceptor Ub on Ub-Degron (**Fig. 2a** inset). Ubc2 was first chemically modified with the bifunctional adaptor molecule **1** ^36^ at the catalytic Cys to form molecule **2**. Subsequently, the S-acetamidomethyl (Acm) group on molecule **2** was removed to expose a β-mercaptoethylamine group for NCL with a Ub thioester (Ub-MesNa) ^36^, generating Ubc2∼Ub carrying an additional thiol group (molecule **3**). Finally, a stable intermediate mimic was obtained by the creation of a disulfide bond between the thiol groups of molecule **3** and a Ub- Degron carrying the K48C point mutation (**Fig. 2b, Fig. S2b**). The intermediate mimic was then mixed with Ubr1 in a 1:1.5 molar ratio and incubated on ice for 30 minutes, followed by vitrification and single-particle cryo-EM analysis (**Fig. S2c**). A dataset of 5083 movie stacks was collected and processed (**Fig. S6a**). The final reconstructed map had an overall resolution of 3.6 Å (**Fig. S6b & c**). Model building was performed by first docking the structure of the initiation complex (Ubr1-Ubc2-Ub-Degron), followed by rigid-body adjustment in Chimera and manual adjustment in COOT ^26^. The final model was refined in real space using PHENIX ^27^ (**Table S2**).

**Figure 2:**
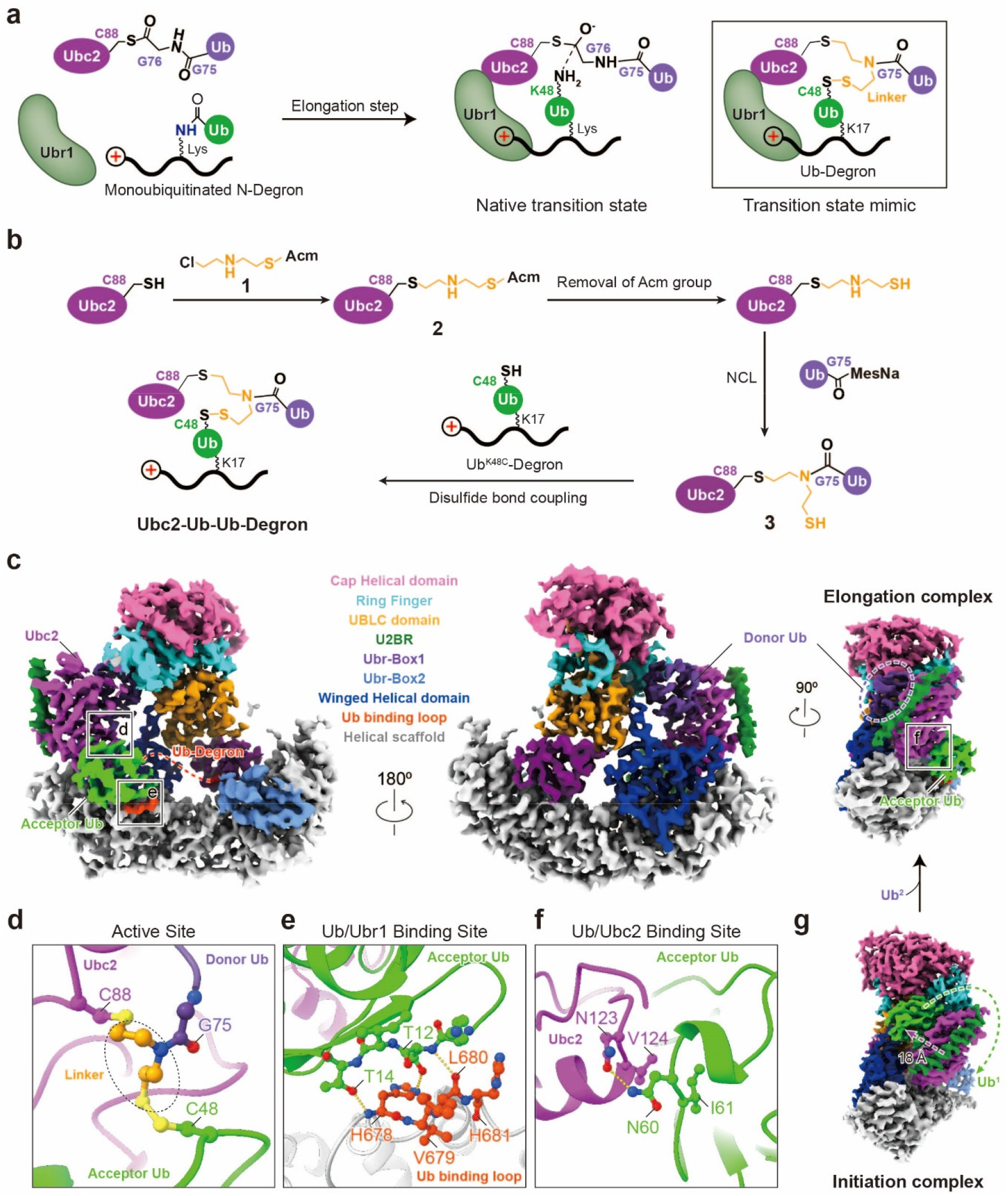
Structure of the elongation complex. **a**, A schematic representation of the transition state of the elongation step. The side chain of K48 on Ub-Degron attacks the thioester bond of Ubc2∼Ub. The inset shows the designed intermediate structure mimicking the transition state of the elongation step. **b**, The synthetic route of the intermediate structure mimicking the transition state of the elongation step. **c**, Cryo-EM maps of the elongation complex (sharpened using a B factor of −96.7 Å^2^, contour level: 0.022). The colour code of Ubr1 is the same as that in Fig. 1b. **d,** Molecular structure of the active site. **e,** Molecular interactions between acceptor Ub and the Ub binding loop of Ubr1. Hydrogen bonds are represented by dotted lines. **f,** Molecular interactions between acceptor Ub and Ubc2. Hydrogen bonds are represented by dotted lines. **g**, A side view of the initiation complex in the same orientation as that of the elongation complex in panel **c**. Displacement of Ubc2, U2BR, and Ub are noted with arrows.

The overall structure of the elongation complex, named Ubr1-Ubc2-Ub-Ub-Degron, adopted a sailboat-like shape similar to that of Ubr1-Ubc2-Ub-Degron (**Fig. 2c**). The linker molecule covalently linked to C88 of Ubc2 forms a disulfide bond with C48 of acceptor Ub and a peptide bond with G75 of donor Ub, as designed (**Fig. 2d**). The acceptor Ub binds to the helical scaffold of Ubr1 at a loop (678-681) located in region C (**Fig. 2e**), which was disordered in the initiation complex (**Fig. 1c**). The acceptor Ub further participated in the recruitment of Ubc2∼Ub by binding at a new interface on Ubc2 (**Fig. 2f**). When mutations were introduced into the Ub binding loop (678-681) on Ubr1 and the new interface (N123 and V124) on Ubc2, the polyubiquitination level on Degron and Ub-Degron were greatly reduced compared to that of the wild-type (**Fig. S7a & b**). Importantly, the Ub binding loop mutant transferred more Ub to Degron than to Ub-Degron, suggesting that the elongation step was impaired (**Fig. S7a**). We further examined the single-turnover ubiquitination of the Ub binding loop mutant using Ubc2 charged with either wild-type Ub or mutant Ub with all lysines mutated to arginines (**Fig. S7c**). Much higher Ub discharge was observed for Degron than Ub-Degron. These results suggested a crucial role of the Ub binding loop on Ubr1 in the elongation step.

In addition to the new interacting surfaces on Ubr1 and Ubc2, U2BR and Ubc2 (including the donor Ub) underwent a displacement of approximately 18 Å, whereas the relative positions of other domains on Ubr1 remained unchanged (**Fig. 2g and Fig. S7d & e**). This displacement of U2BR and Ubc2 repositioned the presumed thioester bond between Ubc2 and the donor Ub so that this bond was approachable by the K48 of the acceptor Ub on Ub-Degron (**Fig. 2d & g**). Together, the two complex structures suggested that the displacement of U2BR on Ubr1 is the key to accommodating extra Ub molecules during the transition from the initiation step to the elongation step.

We further investigated the extensive binding interface between U2BR and Ubc2 which remains the same in both initiation and elongation complex (823.3 Å^2^, **Fig. 3a & b**). Mutations of interface residues either on U2BR (F1190A, Q1186A, F1183A, H1175A) or Ubc2 (L29A, P30A, N37A, W149A) severely impacted the activity of Ubr1 (**Fig. 3f**). Almost no ubiquitination of Degron was observed when including both mutants. We further synthesized a U2BR peptide (Ubr1 1165-1200) and performed an in vitro ubiquitination reaction in the presence of the free U2BR peptide, and dose-dependent inhibition of polyubiquitination was observed (**Fig. 3g**). Using isothermal titration calorimetry (ITC), we quantified the affinity between the U2BR peptide and Ubc2. The dissociation constant (*Kd*) was 143 (±45) nM (**Fig. S8a**). The effect of E1- dependent thioester bond (Ubc2∼Ub) formation in the presence of the U2BR peptide was also examined. Interestingly, an inhibitory effect (IC50 = 9.75 ± 4.34 μM) was observed (**Fig. S8b & c**), which was different from the activation effects of U7BR on Ubc7 ^35^ and the rate-decreasing effects of G2BR on UBE2G2 ^34^. We further tested the accessibility of the catalytic cysteine (Cys88) of Ubc2 using a bulky fluorescent alkylation reagent (BFAR), fluorescein-5-maleimide, in the presence of the U2BR peptide. The results suggested that the U2BR peptide did not enhance the accessibility of this cysteine residue (**Fig. 3h**).

**Figure 3:**
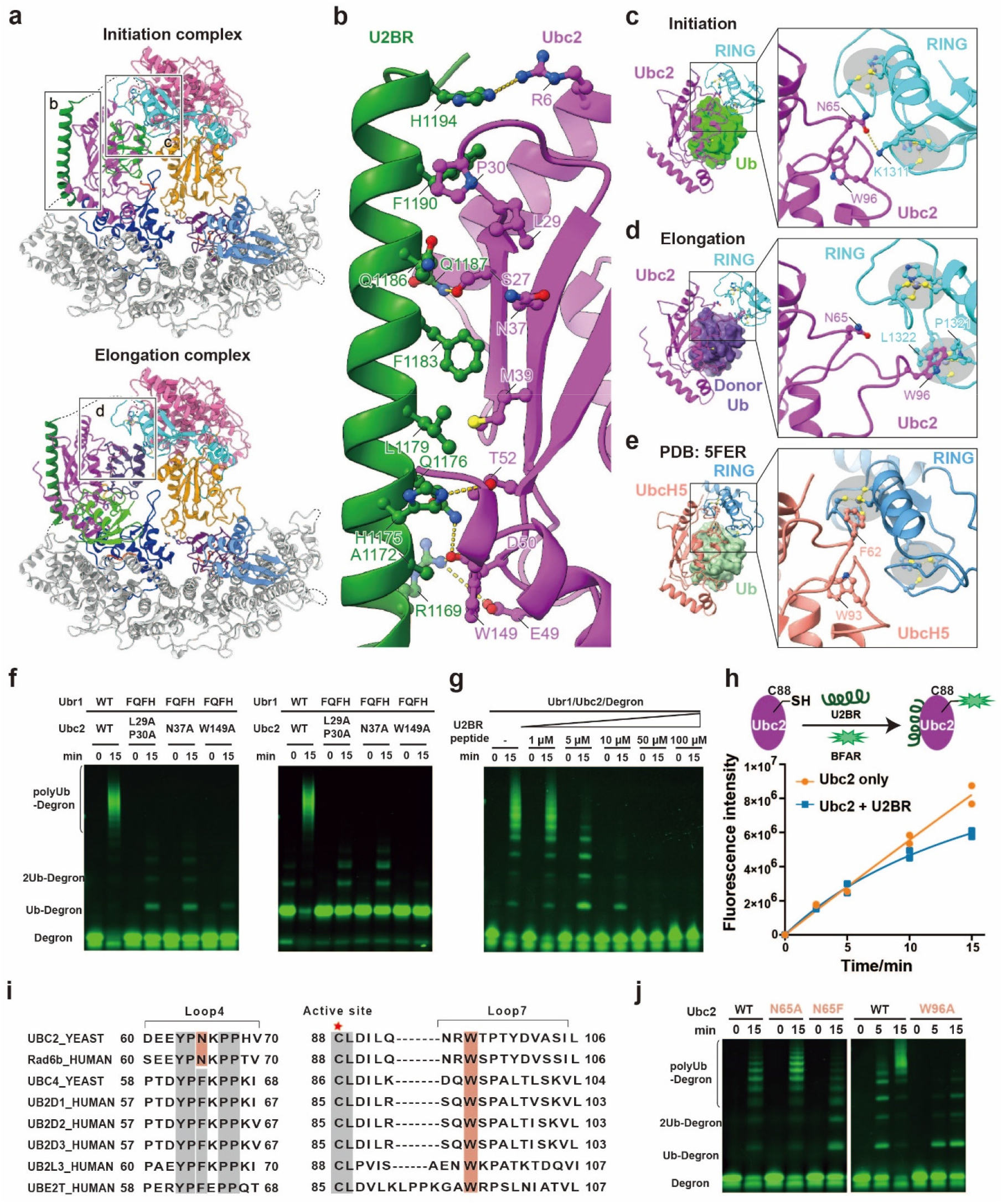
Analysis of the interactions between Ubc2 and Ubr1. **a**, Molecular structures of the initiation complex and the elongation complex. **b**, A close-up view of the interface between Ubc2 and the U2BR of Ubr1. Labelled residues are involved in extensive noncovalent interactions. Hydrogen bonds and electrostatic interactions are represented by dotted lines. **c-e**, Close-up views of the interface between E2 and RING finger domains. **c,** Ubc2 and the RING finger domain in the initiation complex. Hydrogen bonds are represented by dotted lines. **d,** Ubc2 and the RING finger domain in the elongation complex. **e,** UbcH5 and the RING finger domain of TRIM25 (PDB: 5FER) ^38^. **f-g**, In vitro Ubr1-dependent ubiquitination assays. **f,** Ubr1 and Ubc2 mutants at the interface shown in panel **b** were tested. **g**, The inhibition of Ubr1-dependent ubiquitination in the presence of increasing concentrations of a synthetic U2BR peptide. **h**, The accessibility of the catalytic cysteine (Cys88) of Ubc2 tested using a BFAR in the presence or absence of the synthetic U2BR peptide. The average fluorescence from two independent experiments was plotted. **i**, Sequence alignment of multiple E2 enzymes, including yeast and human Ubc2 (also known as Rad6b). Two regions involved in the interaction with the RING finger domain are shown. **j**, In vitro Ubr1-dependent ubiquitination assays investigating the role of N65 and W96 of Ubc2 in the interaction with the RING finger domain of Ubr1.

In both the initiation and elongation structures, we observed smaller interfaces between Ubc2 and the RING finger domain of Ubr1 (410.3 Å^2^ and 208.2 Å^2^, respectively) than were observed in previous studies, such as the interface between UbcH5 (E2) and the RING domain of TRIM25 (547.4 Å^2^, PDB: 5FER, **Fig. 3c-e**) ^37^, although the so-called closed conformation of the E2 and Ub subcomplexes remained the same. Indeed, mapping of conserved E2 residues involved in the RING-E2 interaction showed that Ubc2 does not have the highly conserved aromatic Phe residue involved in the interface, which has been shown to be important for activity ^38^. In yeast and human Ubc2, this residue is mutated to asparagine (**Fig. 3i**). As expected, the N65F mutation of Ubc2 decreased ubiquitination activity in vitro (**Fig. 3j**), suggesting that this Phe residue is not required for the interaction between Ubc2 and the RING finger domain of Ubr1. Interestingly, the N65A mutation increased the amount of polyubiquitinated Degron (**Fig. 3j**), suggesting that N65 may not be important for the elongation of Ubr1-mediated ubiquitination. Notably, the interface between Ubc2 and the RING domain changed in the elongation structure. A conserved tryptophan residue (W96 of Ubc2) flipped out and interacted with the RING domain instead of N65 (**Fig. 3d & i**). The W96A mutation did not affect the initiation step but severely decreased the amount of polyubiquitinated Degron (**Fig. 3j**). This result suggested that the new interface involving the tryptophan residue is critical for the elongation of Ub chain and may play a role in other E2-E3 systems.

## DISCUSSION

Visualizing E3-mediated substrate ubiquitination is of great importance ^39^, and it is also informative for the development of novel therapeutic strategies such as PROTAC ^16,17^. The chemical trapping of ubiquitinated intermediates has played critical roles in the mechanistic understanding of various E3 ligases. The Lima group engineered an E2 to trap the E3_Siz1_/E2_Ubc9_- SUMO/PCNA complex, demonstrating that E3 could bypass E2 specificity to force-feed a substrate lysine into the E2 active site ^40^. The Schulman group designed a chemically trapped complex of neddylated CRL1^β-TRCP^-UBE2D-Ub-phosphorylated IκBα, showing that the E3 ligase CRL1^β-TRCP^ primed and positioned Ub and the substrate lysine for transfer ^38^. Here, through the chemical synthesis of two ubiquitination intermediate mimics representing the initiation and elongation steps and single-particle cryo-EM analysis, we revealed the mechanism of Ubr1-mediated ubiquitination involved in the N-end rule pathway. Our cryo-EM structures suggested the rapid and linkage-specific ubiquitination of Ubr1 (**Fig. 4**). Specifically, substrates bearing destabilizing amino-terminal degrons are captured by Ubr1 through the Ubr-Box. Ub-charged E2, Ubc2∼Ub, is recruited through the U2BR and the RING domain. The positions of these domains on Ubr1 facilitate Ub thioester transfer to a Lys residue near the degron (initiation step). The helical scaffold of Ubr1 provides an additional anchor (the Ub binding loop) for Ub on the newly formed monoubiquitinated substrate. Ub also participates in the rearrangement of U2BR-Ubc2 by interacting with a new interface on Ubc2. The specific binding between the Ub binding loop and Ub ensures close proximity of K48 on Ub and the newly formed thioester bond at the active site of Ubc2, which facilitates the transfer of the second Ub (elongation step). We further speculate that similar rearrangements occur for subsequent elongation steps. The most distal Ub is always engaged by the Ub binding loop on Ubr1 to ensure linkage specificity of the polyubiquitin chain (**Fig. 4**).

**Figure 4:**
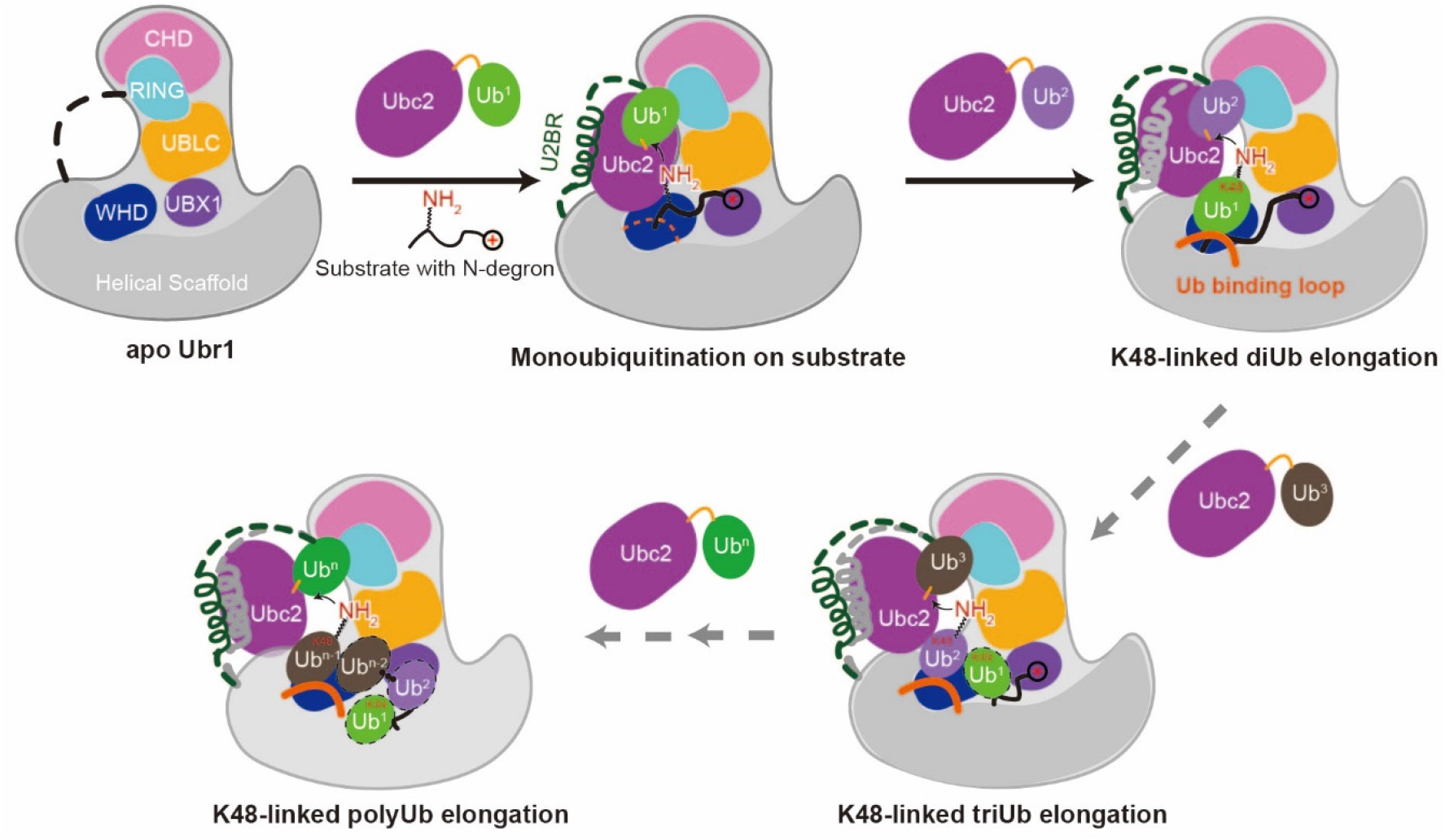
Model of Ubr1-mediated polyubiquitination. A cartoon representation of Ubr1-mediated polyubiquitination starting from a degron peptide with a positively charged N-terminal residue. The first two steps correspond to the initiation and elongation structures described in this study. Subsequent elongation of the polyubiquitin chain is hypothetical. The Ub molecules being conjugated are sequentially numbered as Ub^1^, Ub^2^, …, Ub^n^.

Our structures revealed that U2BR plays a key role in the recruitment of Ubc2. Notably, single-particle analysis of Ubr1 alone showed that the cap helical domain was very flexible without Ubc2 and the substrate (**Fig. S9**). U2BR could not be resolved from 2D class averages and 3D reconstruction, suggesting that the chemical synthesis of intermediate mimics is the key to stabilizing Ubr1, which leads to structure determination at near-atomic resolution. The strategies presented in this study can be adopted to investigate the mechanism of other E3 ligases.

In humans, Johanson-Blizzard syndrome is a rare and severe autosomal recessive genetic disorder caused by mutations to Ubr1 ^9^. Human Ubr1 shares 20% sequence identity with the yeast homologue, especially in Ubr-Box1, Ubr-Box2, region C of the helical scaffold, the U2BR and the RING finger domain. Given the similarity, human Ubr1 is very likely to have a similar mechanism. Our structures provide a molecular basis to understand the pathogenesis of this disease.

## ACKNOWLEDGEMENTS

We thank the staff at the National Cryo-Electron Microscopy Facility at the Frederick National Laboratory and the Advanced Electron Microscopy Facility at the University of Chicago for the help in cryo-EM data collection. Funding: Funding for this work was, in part, provided by the Catalyst Award from the Chicago Biomedical Consortium. This work was supported by Chicago Biomedical Consortium Catalyst Award C-086 to M.Z. We thank the National Key R&D Program of China (No. 2017YFA0505200), NSFC (91753205) for financial support. This research was, in part, supported by the National Cancer Institute’s National Cryo-EM Facility at the Frederick National Laboratory for Cancer Research under contract HSSN261200800001E. This material is based upon work supported by the National Science Foundation under Grant No. 2030381, the graduate research award of Computing and Software Systems division, and the start-up fund 74–0525 at University of Washington Bothell to D.S.. Any opinions, findings, and conclusions or recommendations expressed in this material are those of the authors and do not necessarily reflect the views of the National Science Foundation. Molecular graphics and analyses were performed with UCSF ChimeraX, developed by the Resource for Biocomputing, Visualization, and Informatics at the University of California, San Francisco, with support from National Institutes of Health R01-GM129325 and the Office of Cyber Infrastructure and Computational Biology, National Institute of Allergy and Infectious Diseases.

## AUTHOR CONTRIBUTIONS

M.P., M.Z., L.L. and Y.Y. designed all the experiments and interpreted the results. M.P., Q.Z. and L. L. designed the synthetic route for chemically synthesized ubiquitination initiation and elongation intermediate mimics. T.W. synthesized the fluorescently labelled Ub-Degron and the elongation intermediate mimic. L.J.L. synthesized the fluorescently labelled Degron and the initiation intermediate mimic. M.P., Y.Y., D.S., and M.Z. performed cryo-EM data collection and processing. J.M. performed the in vitro ubiquitination assays with Ubr1 and Ubc2 mutants. Q.Z. performed characterization of the U2BR peptide on the enzymatic properties of Ubc2. T.W., Y.Y., R.D., J.M., H.A. and Y.X. cloned, expressed, and purified Ubr1, Ubc2 and their mutants. M.Z., M.P., and L.L. wrote the paper. M.Z., L.L., Y.Y. and M.P. supervised the project.

## Supplementary Figures and Legends

**Fig. S1:**
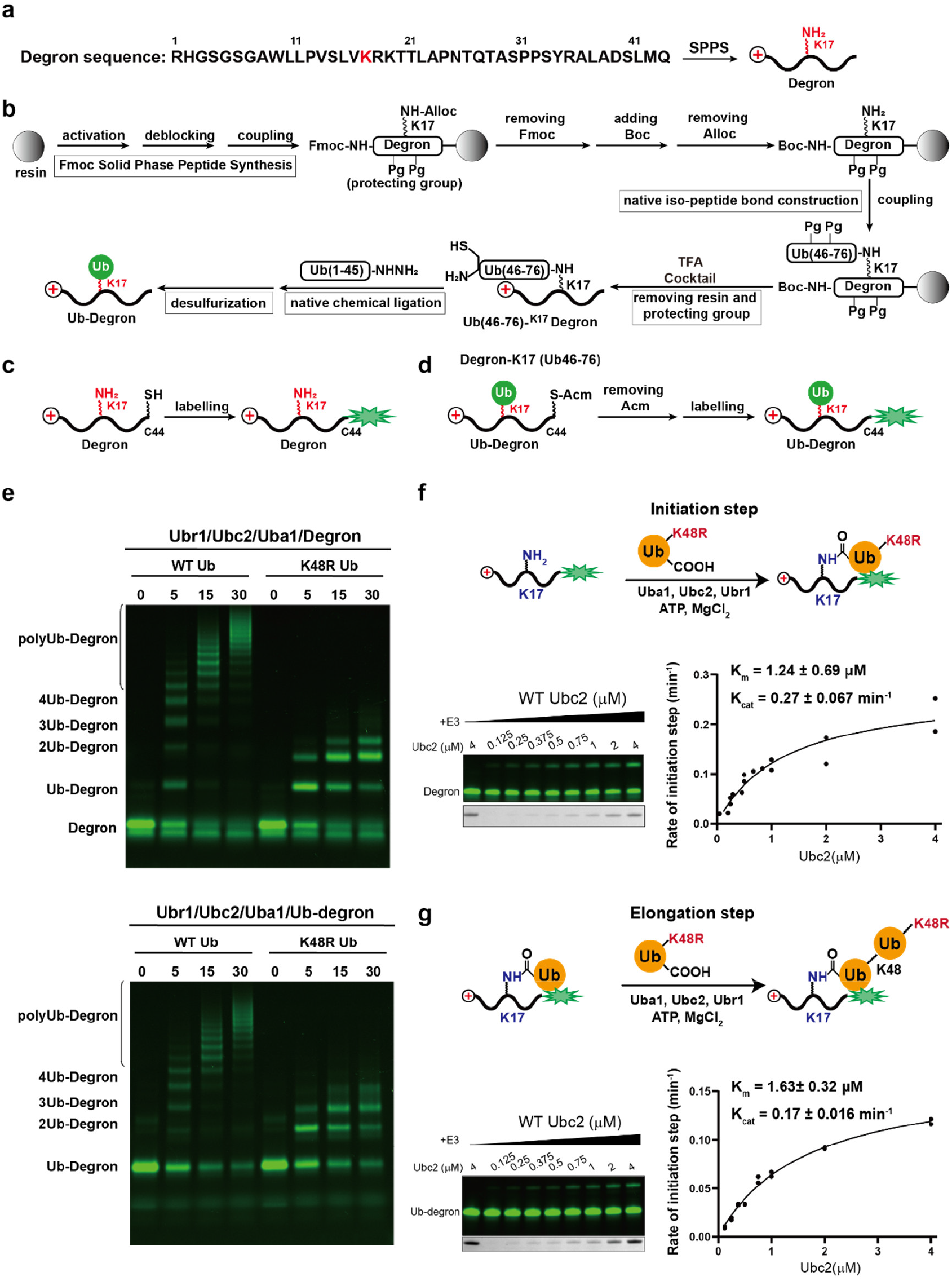
Ubr1-mediated K48-linked polyubiquitination of degron peptides. **a**, Amino acid sequence of the degron peptide (Degron). **b**, The synthetic route of the monoubiquitinated degron peptide (Ub-Degron). **c-d**, Fluorescent labelling of Degron (**c**) and Ub-Degron (**d**). **e**, In vitro Ubr1-dependent ubiquitination assays using fluorescent Degron (top) and Ub-Degron (bottom) as substrates. **f-g**, Quantitative evaluation of the Ubr1 enzyme kinetics for ubiquitination initiation (**f**) and the first step of elongation (**g**). Averages of two independent experiments were plotted and fit to the Michaelis–Menten model to estimate the *K_m_* and *K_cat_*.

**Fig. S2:**
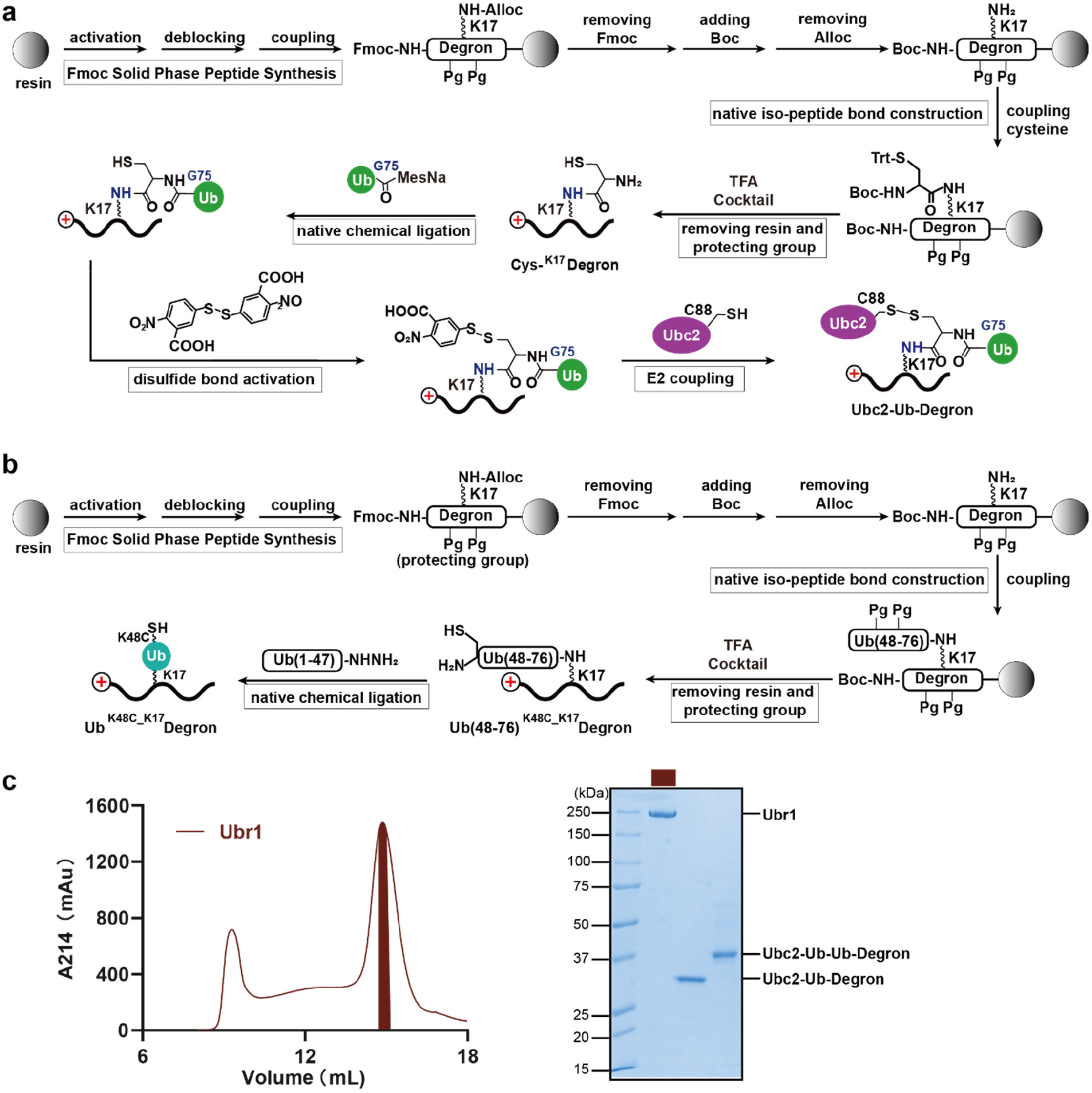
Synthesis and purification of the intermediate structures. **a**, The synthetic route of the designed structure mimicking the transition state of the initiation step. **b**, The synthetic route of Ub(K48C)-^K17^Degron. **c**, A gel filtration chromatogram of Ubr1 (left) and an SDS-PAGE gel of purified Ubr1 and designed stable intermediate structures Ubc2- Ub-Degron and Ubc2-Ub-Ub-Degron (right).

**Fig. S3:**
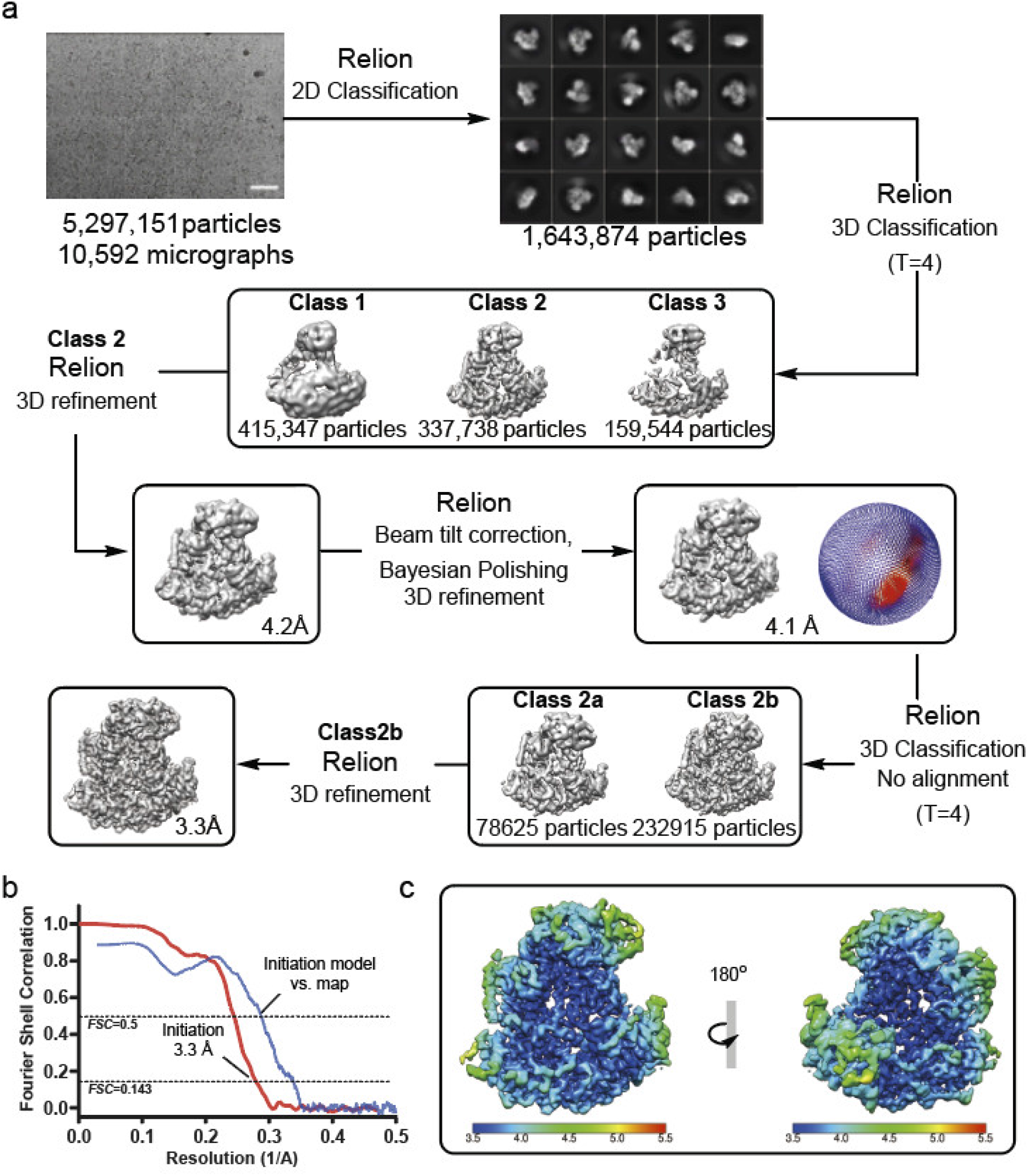
Single-particle cryo-EM analyses of the Ubr1-Ubc2-Ub-Degron dataset. **a,** The workflow of data processing. The dataset was subjected to particle selection, 2D classification, and multiple rounds of 3D classification. A representative micrograph (scale bar corresponds to 50 nm) and representative 2D class averages are shown. The distribution of the Euler angles is shown next to the map. **b,** Fourier shell correlation (FSC) curve of the masked map after Relion postprocessing. The resolution was determined by the FSC=0.143 criterion. The model vs. map FSC curve is also shown. **c,** Local resolution of the map calculated using Relion.

**Fig. S4:**
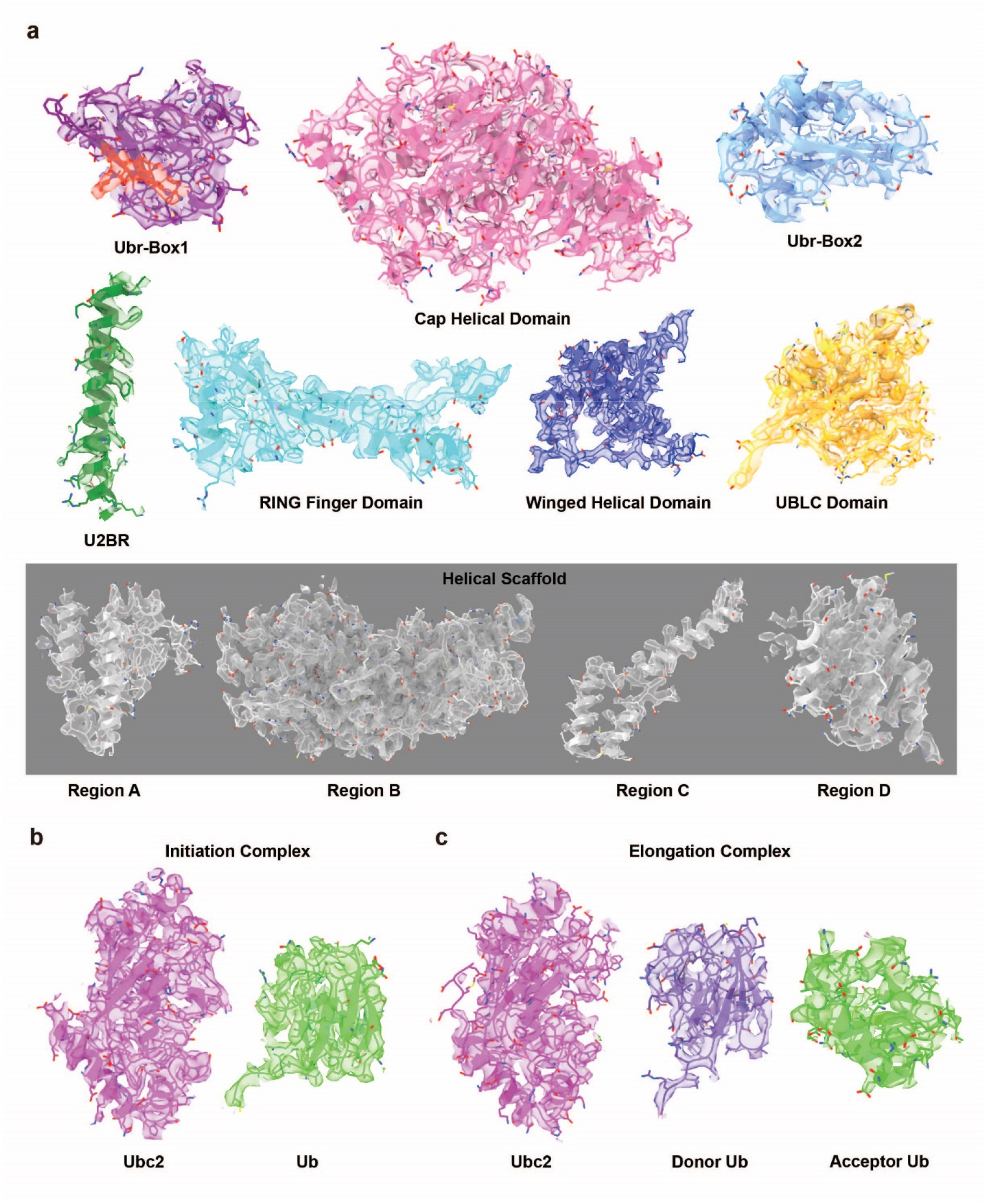
Representative cryo-EM densities of the initiation and elongation complexes. **a**, Individual domains in the initiation complex. **b**, Ubc2 and Ub in the initiation complex. **c**, Ubc2, donor Ub and acceptor Ub in the elongation complex. Maps in panels **a** and **b** were sharpened using a B factor of −96.5 Å^2^ and contoured at a level of 0.03. Maps in panel **c** were sharpened using a B factor of −96.7 Å^2^ and contoured at a level of 0.022.

**Fig. S5:**
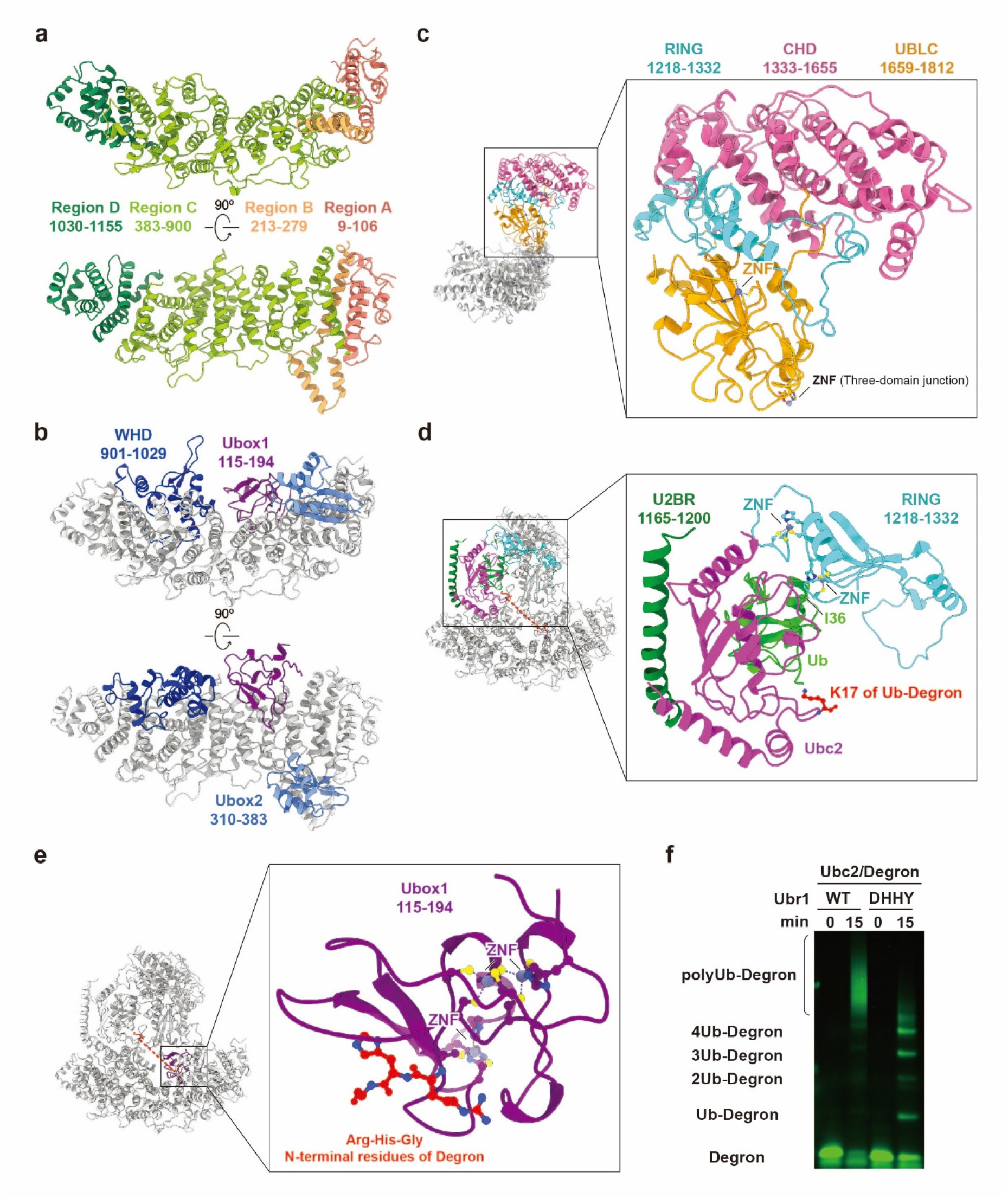
Molecular structures of Ubr1 domains. **a,** The helical scaffold of Ubr1 consists of four separate regions. **b,** The three domains located around the helical scaffold, Ubr-Box1 (Ubox1, purple), Ubr-Box2 (Ubox2, light blue) and winged helical domain (WHD, dark blue). **c,** The three domains above the helical scaffold, the RING finger domain (cyan), cap helical domain (pink) and UBLC domain (yellow). The RING finger domain joins the cap helical domain and UBLC domain. An additional zinc finger motif (ZNF) in UBLC is labelled. **d,** The U2BR (forest green), Ubc2 (magenta)∼Ub (lime) and the RING finger domain form the catalytic module of Ubr1. Two zinc finger motifs (ZNF) in the RING finger domain are labelled. **e,** A close-up view of substrate-engaged Ubr-Box1. Three zinc finger motifs (ZNF) are labelled. **f,** In vitro Ubr1-dependent ubiquitination assay. A quadruple mutant (H161A, Y933A, D1175A, and H1763A) of the residues involved in the interface between Ubox1, WHD, and UBLC (three-domain junction, shown in **Fig. 1f**) was tested.

**Fig. S6:**
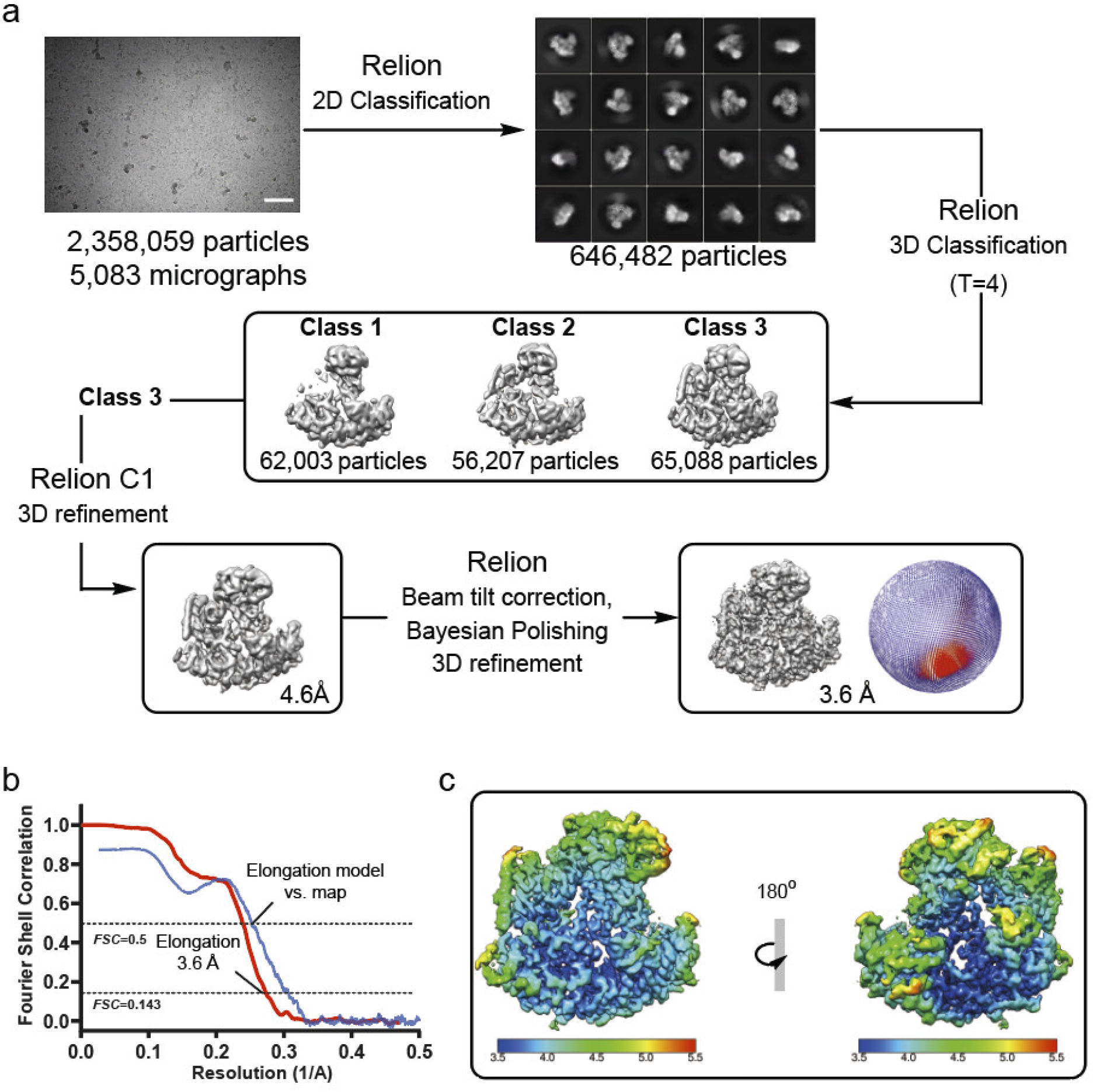
Single-particle cryo-EM analyses of the Ubr1-Ubc2-Ub-Ub-Degron dataset. **a,** The workflow of data processing. The dataset was subjected to particle selection, 2D classification, and multiple rounds of 3D classification. A representative micrograph (scale bar corresponds to 50 nm) and representative 2D class averages are shown. The distribution of the Euler angles is shown next to the map. **b,** Fourier shell correlation (FSC) curve of the masked map after Relion postprocessing. The resolution was determined by the FSC=0.143 criterion. The model vs. map FSC curve is also shown. **C,** Local resolution of the map calculated using Relion.

**Fig. S7:**
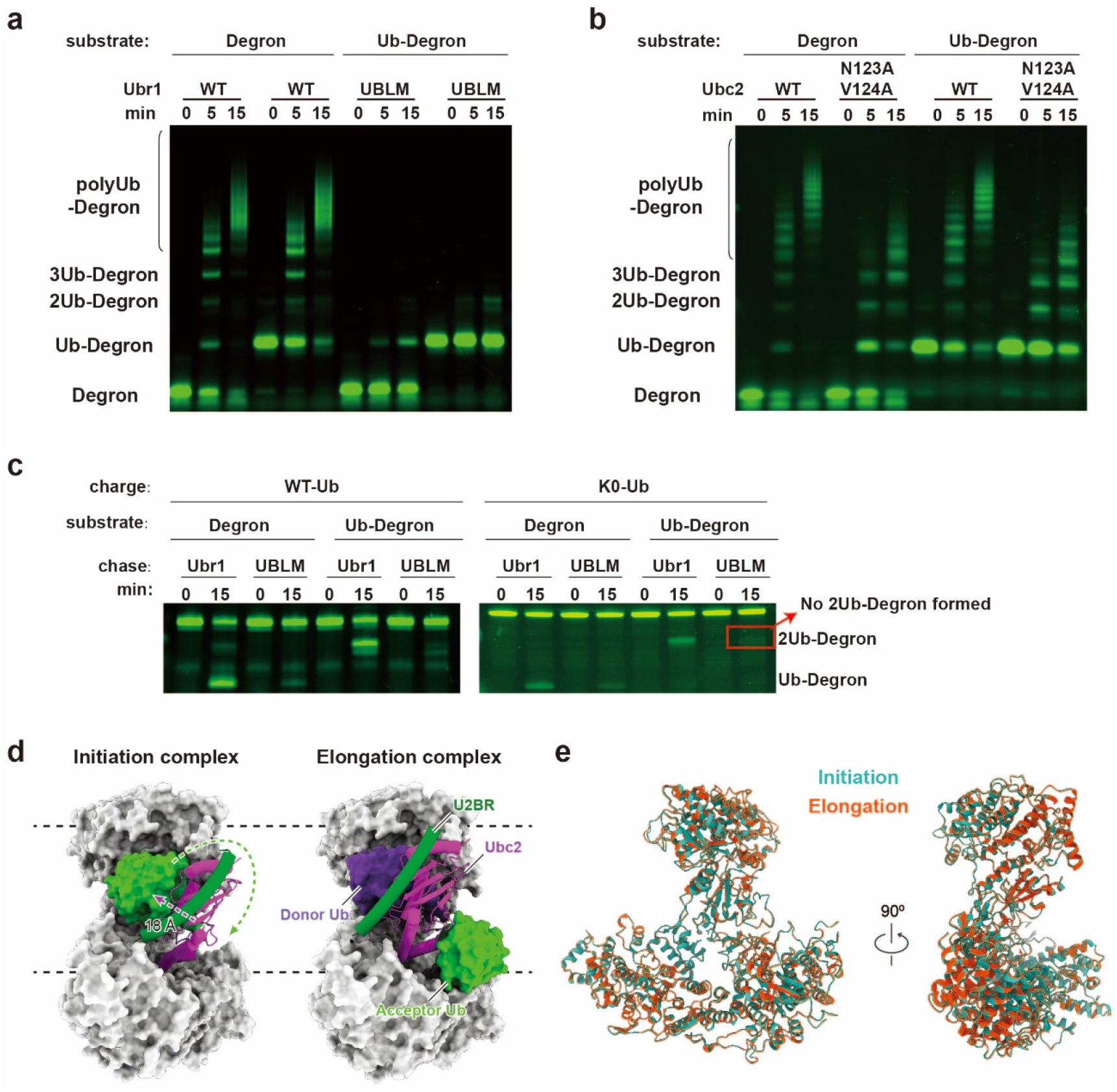
Characterization of the acceptor Ub interactions in the elongation complex. **a-b**, In vitro Ubr1-dependent ubiquitination assays. Mutations of the Ub binding loop (H678A/V679A/L680A/H681A, UBLM, panel **a**) and the Ubc2 interface (N123A/V124A, panel **b**) were tested. **c**, Single-turnover ubiquitination assay of wild-type Ubr1 and the Ubr1 mutant (UBLM) using Ubc2 charged with either wild-type Ub or K0-Ub (all lysines mutated to arginines). The red box highlights that the Ubr1 mutant (UBLM) could not mediate Ub thioester discharge from Ubc2∼Ub to Ub-degron (a defect in elongation). **d**, Side views of the initiation and elongation complexes showing the displacement of U2BR, Ubc2, and Ub. **e**, Alignment of Ubr1 in the initiation and elongation complexes.

**Fig. S8:**
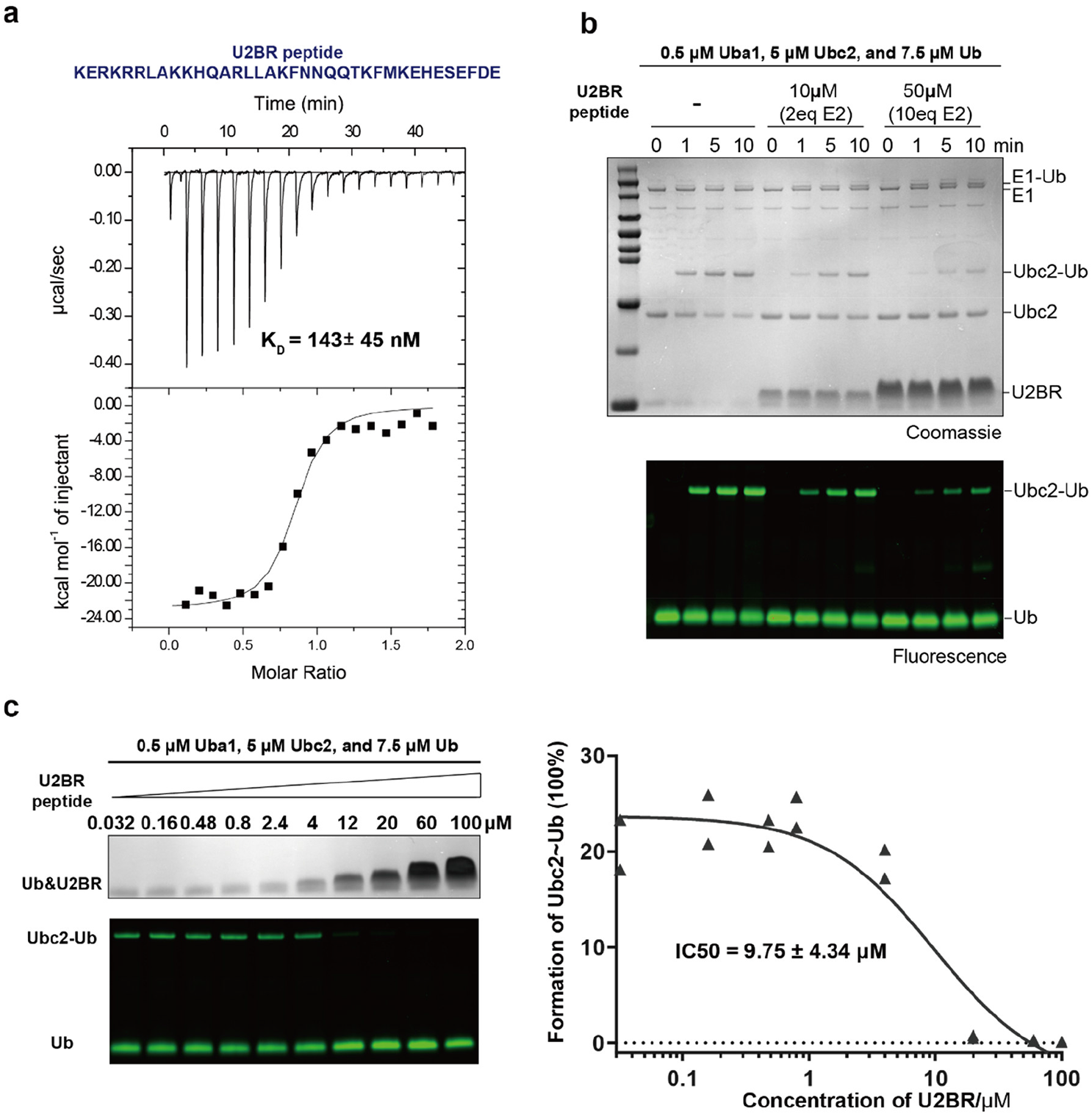
Characterization of the U2BR peptide on the enzymatic properties of Ubc2. **a**, ITC measurement of the binding between Ubc2 and a synthetic U2BR peptide. **b**, E1- dependent Ubc2∼Ub thioester formation in the presence of the synthetic U2BR peptide. **c**, Quantitative evaluation of the inhibitory effects of the synthetic U2BR peptide on E1-dependent Ubc2∼Ub thioester formation. Averages of two independent experiments were plotted and fit to estimate the IC50 of the synthetic U2BR peptide.

**Fig. S9:**
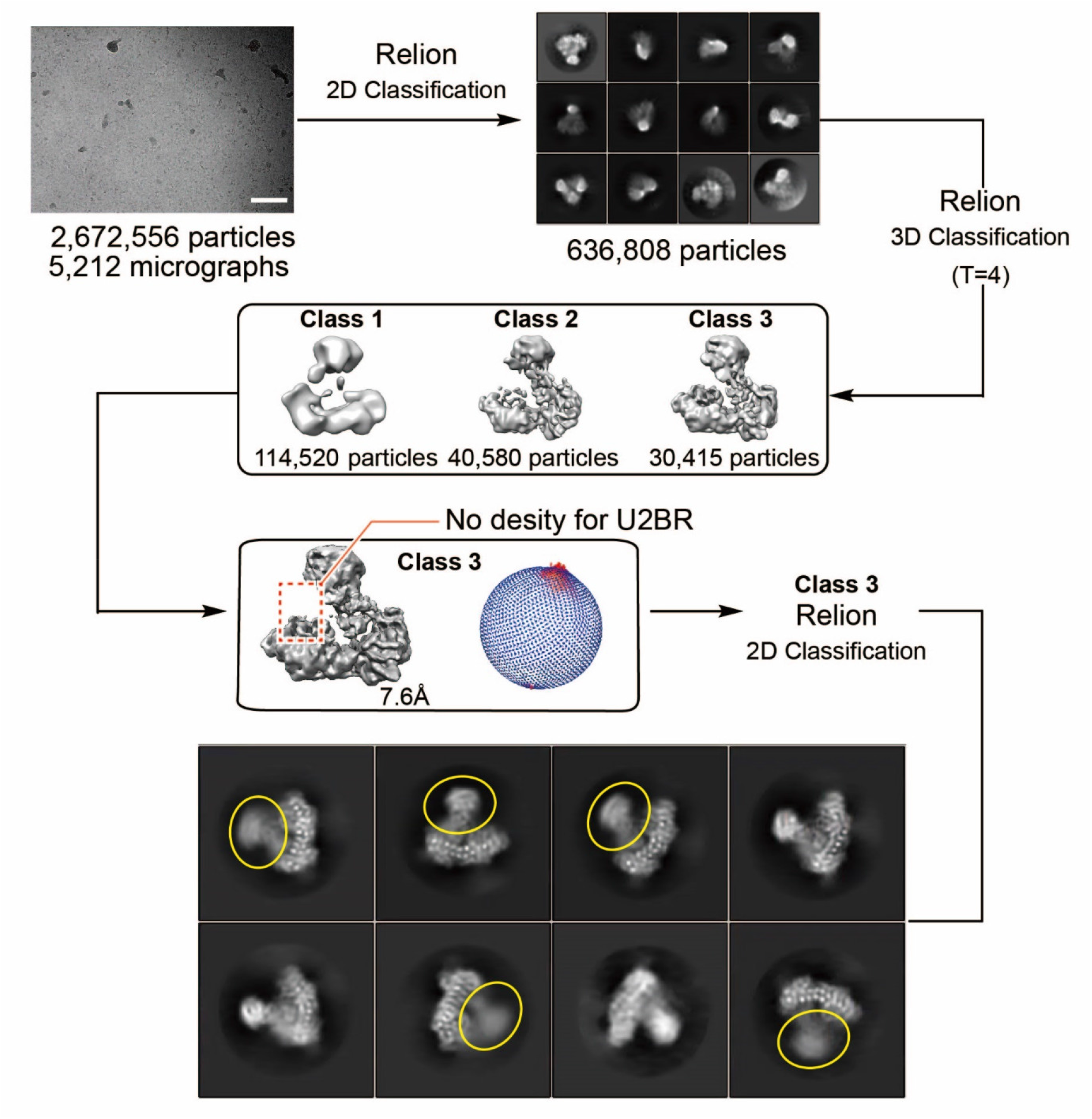
Single-particle cryo-EM analyses of Ubr1. The dataset was subjected to particle selection, 2D classification, and multiple rounds of 3D classification. A representative micrograph (scale bar corresponds to 50 nm) and representative 2D class averages are shown. Flexible cap helical domains in the 2D class averages are highlighted in yellow circles. The distribution of the Euler angles is shown next to the map.

**Table S1:** Statistics of cryo-EM data collection and processing.

|  | **Initiation complex** | **Elongation complex** | **Apo Ubr1** |
| --- | --- | --- | --- |
| Detergent | FOM (0.001%) | FOM (0.001%) | FOM (0.001%) |
| Microscope | Krios (UChicago) | Krios (UChicago) | Krios (NCI) |
| Magnification | 81,000 | 81,000 | 81,000 |
| Voltage (kV) | 300 | 300 | 300 |
| Spherical aberration (mm) | 2.7 | 2.7 | 2.7 |
| Detector | K3 | K3 | K3 |
| Camera mode | Super resolution counting | Super resolution counting | Super resolution counting |
| Exposure rate (e<sup>-</sup>/pixel/s) | 15 | 15 | 15 |
| Total exposure (e<sup>-</sup>/Å<sup>2</sup>) | 50 | 50 | 50 |
| Defocus range (μm) | -1.0 to -2.5 | -1.0 to -2.5 | -1.0 to -2.5 |
| Pixel size (Å) | 1.063 | 1.063 | 1.122 |
| Mode of data collection | Image shift | Image shift | Image shift |
| Energy filter | 20 eV slit | 20 eV slit | 20 eV slit |
| Software for data collection | EPU | EPU | Latitude S |
| Number of micrographs | 10,592 | 5,083 | 5,212 |
| Symmetry imposed | C1 | C1 | C1 |
| Box size (pixel) | 320 | 320 | 320 |
| Initial particle images (no.) | 5,297,151 | 2,358,059 | 2,672,556 |
| Particle images for 3D (no.) | 1,643,874 | 646,482 | 636,808 |
| Final particle images (no.) | 232,915 | 65,088 | 30,415 |
| Map resolution, masked (Å) | 3.35 | 3.67 | 4.56 |
| B-factor estimated (Å<sup>2</sup>) | 96.5 | 96.7 | 119.0 |
| EMD accession code | 23806 | 23807 | - |

**Table S2:** Statistics of cryo-EM model refinement and geometry.

|  | Initiation complex | Elongation complex |
| --- | --- | --- |
| Composition (#) |  |  |
| Chains | 5 | 7 |
| Atoms | 15934 | 16488 |
| Residues | 1967 | 2036 |
| Water | 0 | 0 |
| Ligands | ZN: 7 | ZN: 7 |
|  |  | ETE: 1 (chemical linker) |
| Bonds (RMSD) |  |  |
| Length (Å) | 0.003 | 0.004 |
| Angles (°) | 0.604 | 0.646 |
| MolProbity score | 1.74 | 1.77 |
| Clash score | 10.39 | 11.19 |
| Ramachandran plot (%) |  |  |
| Outliers | 0.00 | 0.00 |
| Allowed | 3.29 | 3.27 |
| Favored | 96.71 | 96.73 |
| Rotamer outliers (%) | 0.28 | 0.27 |
| Cβ outliers (%) | 0.00 | 0.00 |
| Peptide plane (%) |  |  |
| Cis proline/general | 0.0/0.0 | 0.0/0.0 |
| Twisted proline/general | 0.0/0.0 | 0.0/0.0 |
| CaBLAM outliers (%) | 1.81 | 1.5 |
| ADP (B-factors) |  |  |
| Iso/Aniso (#) | 15934/0 | 16488/0 |
| min/max/mean |  |  |
| Protein | 12.90/108.84/44.62 | 30.00/164.26/82.92 |
| Ligand | 53.14/105.95/73.23 | 94.39/153.23/112.90 |
| PDB accession code | 7MEX | 7MEY |

## METHODS

### Cloning and plasmid construction

The plasmid containing yeast (*Saccharomyces cerevisiae*) Ubr1 was purchased from Addgene (plasmid # 24506) ^41^. DNA sequence of yeast (*Saccharomyces cerevisiae*) Ubc2 was synthesized and codon optimized for *E. coli* overexpression by Genscript. The gene was further cloned between the NdeI and XhoI sites of the vector pET-28a containing an N-terminal His tag followed by a HRV3C protease cleavage site. Variants of Ubr1 and Ubc2 were generated using site directed mutagenesis. Human Uba1 was cloned in pET-28a vector containing an N-terminal His tag. DNA sequences of wildtype ubiquitin (Ub), Ub mutants including K48R, G76C, K0 (all 7 lysine residues mutated to arginine), and AC-Ub (a Ub variant with additional two amino acids Ala-Cys at the N terminus) were synthesized and codon optimized for *E. coli* overexpression by Genscript. The genes were further cloned between the NdeI and XhoI sites of the vector pET- 22b.

### Protein expression and purification

Wildtype Ub and Ub mutants were purified as previously described ^42^. Briefly, plasmids for overexpression were transformed to *E. coli* BL21(DE3) competent cells. The *E. coli* cells were grown in Luria broth (LB) media containing 50 µg/mL ampicillin until OD600 reached 0.8 and were induced by isopropyl β-D-1-thiogalactopyranoside (IPTG) at a final concentration of 0.4 mM followed by overnight incubation at 25 °C. Cells were pelleted at 4,000 rpm for 30 min at 4 °C, resuspended in ddH2O and lysed by ultra-sonication for 30 min in an ice bath. The cell lysates were supplemented with 1% perchloric acid to precipitate non-relevant proteins which were then cleared using centrifugation (30 min, 15,000×g, 4 °C). Ub and its variants were further purified using a Mono S cation exchange column (GE Healthcare), followed by dialysis into a buffer containing 50 mM HEPES, pH 7.5, and 150mM NaCl. The peak fractions were pooled and concentrated to 20 mg/ml.

Plasmids containing Ubc2 and its variants were transformed to *E. coli* BL21(DE3) competent cells. The *E. coli* cells were grown in LB media containing 20 µg/mL kanamycin until OD600 reached 0.6, and were induced by IPTG at a final concentration of 0.4 mM followed by overnight incubation at 18 °C. Cells were harvested by centrifugation at 4,000 rpm for 30 min and then lysed by sonication in the lysis buffer (50 mM HEPES, pH 7.5, 150mM NaCl, 20mM imidazole and 1mM PMSF). After centrifugation at 12,000 rpm for 30 min, the supernatant was loaded onto a Ni-NTA affinity column. The proteins were eluted with the elution buffer (50 mM HEPES, pH 7.5, 150mM NaCl, and 400mM imidazole), and further purified by a Superdex 200 size-exclusion column (GE Healthcare) equilibrated in a buffer containing 50 mM HEPES, pH 7.5, and 150 mM NaCl.

Yeast Ubr1 and its variants was expressed as previously described ^33^. Briefly, single colony of yeast were grown in SD medium at 30°C until OD600 reached ∼1.0. The cells were pelleted at 5,000 g, washed once with cold phosphate buffer saline (PBS), and then resuspended (6 mL buffer per 1 g of pellet) in the lysis buffer (50 mM HEPES, pH 7.5, 0.15 M NaCl, protease inhibitor cocktail (Roche), 1mM PMSF, and 10% glycerol,). The resuspended yeast cells were dropped into liquid nitrogen, and the frozen pellet balls were ground to fine powder in liquid nitrogen using a cryogenic impact grinder (SPEX™ SamplePrep 6870 Freezer/Mill). The powder was further thawed and centrifuged at 11,200 g at 4°C for 30 min. The supernatant was loaded onto anti-DYKDDDDK (FLAG) affinity resin (Thermo Scientific, #. A36803), followed by extensive wash. Finally, the FLAG-tagged Ubr1 was eluted with 1mg/mL FLAG peptide and further purified by a Superose 6 size-exclusion column (GE Healthcare) equilibrated in a buffer captaining 50 mM HEPES, pH 7.5, and 150mM NaCl.

### Peptide Synthesis

All peptides were synthesized using standard Fmoc solid phase peptide synthesis (SPPS) protocols under standard microwave conditions (CEM Liberty Blue). Fmoc-hydrazine 2- chlorotrityl chloride PS resin and Rink Amide MBHA PS resin were used for peptides synthesis. The coupling cycle was programed as previously reported ^18^. Briefly, 10% piperidine in DMF with 0.1 M Oxyma (1 min at 90 °C) was applied as deprotection condition, and 4-fold of 0.2 M Fmoc-protected amino acid, 1.0 M DIC, and 1.0 M Oxyma in DMF (10 min at 50 °C for His and Cys, 90 °C for other residues were applied as amino acid coupling condition. Specifically, in peptide Ub(46-76)-^K17^Degron-NH_2_ and Ub(48-76)-^K17^Degron-NH_2_, Fmoc-Lys (Alloc)-OH was coupled at position 17 for the orthogonal protection. When the backbone coupling was finished, the Alloc protecting group was removed by [(Ph_3_)P]_4_Pd/Ph_3_SiH as previously described ^43^, and then the ε-amino group on Lys17 can be further coupled with successive sequence (Ub48-76) or (Ub46-76). After the completion of SPPS, the resulting peptide-resin was cleaved in cleavage cocktail (87% trifluoroacetic acid, 5% water, 5% thioanisole, 3% 1,2-ethanedithiol) for 2 h at 25 °C. Crude peptides were precipitated with cold diethyl ether, analyzed and purified by reversed-phase high-performance liquid chromatography (RP-HPLC).

### Preparation of fluorescence labeled Degron and Ub-Degron

Degron was direct obtained from SPPS as described above. K17-linked mono ubiquitinated degron (Ub-^K17^Degron) was synthesized from two fragments, Ub(1-45)-NHNH_2_ and Ub(46-76)- ^K17^Degron-NH_2_. Ala46 in the latter fragment was temporally mutated to Cys to enable native chemical ligation of these two fragments. After ligation the thiol group of Cys46 was removed through desulfurization reaction to produce the native Ala46 as shown in Fig S1b. For fluorescence labeling of Degron, we introduced an additional Cys at the C-terminus of Degron to enable site-specific labeling. In the Ub-Degron case, a Cys(Acm) was introduced in the C-terminus of Degron to orthogonally protect this thiol group from being desulfurized. After purification, the Acm group was removed from the obtained Ub-Degron-Cys(Acm) to free the thiol group for labeling. For labeling reaction, 2 mg lyophilized dry power of Degron-Cys or Ub- Degron-Cys was dissolved in 50 mM HEPES, 150mM NaCl, pH 7.5. Then 2 equivalents of fluorescein-5-maleimide (Invitrogen, #F150) was added and the mixture was incubated at room temperature for 20 min, followed by buffer exchange in (50 mM HEPES, 150mM NaCl, pH 7.5) using a Superdex peptide size-exclusion column (GE Healthcare) to give the fluorescence labeled degron and Ub-degron.

### Ubiquitination assay with fluorescent Degron or Ub-Degron

In vitro ubiquitination assays were performed with 0.1 μM Uba1, 4 μM Ubc2, 0.25 μM Ubr1, 5 μM fluorescent Degron or Ub-Degron, and 80 μM Ub at 30 °C in the reaction buffer (50 mM HEPES pH 7.5, 0.15 M NaCl, 10 mM MgCl_2_, and 5 mM ATP). The reactions were terminated by adding 4 × SDS-sample buffer, followed by SDS-PAGE. Unless indicated otherwise, same concentrations of Ubr1 and Ubc2 variants as the respective wildtype were used in the assay.

### Single-turnover measurement of Ub transfer in the initiation and elongation steps

To monitor the single-turnover of Ub transfer in the initiation step, i.e., Ubr1 transfers Ubc2∼Ub to Degron, a pulse-chase experiment that eliminates the effects of UBA1-dependent formation of Ubc2∼Ub intermediate was performed. The pulse reaction generated a thioester-linked Ubc2∼Ub intermediate with 5 μM Ubc2, 7.5 μM fluorescent Ub, 0.5 μM UBA1 in a buffer containing 50 mM HEPES pH 8.0, 150 mM NaCl, 10 mM MgCl_2_, and 10 mM ATP. The reaction mixture was incubated at room temperature for 10 min, and quenched with 50 mM EDTA on ice for 5 min. A final concentration of 0.5 μM Ubr1 and 25 μM fluorescently labeled Degron was added for the chase reaction which was then incubated at 30°C for 5 minutes. The reaction was stopped by in 2× SDS-sample buffer (pH < 3), followed by SDS-PAGE. Same concentrations of Ubr1 and Ubc2 variants as the respective wildtype were used. The experimental setup for the elongation step was similar to the initiation step, except that fluorescently labeled Ub-Degron was used instead of Degron.

### Michaelis-Menten constant (Km) measurement of the initiation and elongation steps

For K_m_ measurement of the initiation step, two prepared mixtures are required. Mixture 1 consists of Uba1 and Ub (K48R), and Mixture 2 consists of Ubr1 and fluorescently labelled Degron. Both mitures were prepared in the reaction buffer (50 mM HEPES pH 7.5, 150 mM NaCl, 10 mM MgCl_2_, and 5 mM ATP). Ubc2 was prepared as a twofold dilution series from the stock, and then introduced into the solution containing equal amounts of the 2 mixtures to initiate the reaction. The final concentrations were 80 μM Ub (K48R), 0.1 μM Uba1, 0.25 μM Ubr1 and 5 μM fluorescently labelled Degron. Reactions were quenched after 1 minute at 30°C using 2 × SDS-sample buffer, followed by SDS-PAGE. The gels were imaged on ChemiDoc MP Imaging System. Substrate and product bands were individually quantified as a percentage of the total signal for each time point using ImageLab (Bio-Rad). Ratios of ubiquitinated products relative to the total signal were plotted against the concentration of Ubc2 and fit to the Michaelis–Menten equation to estimate K_m_ in GraphPad Prism 8. The experimental setup for K_m_ measurement of the elongation step was similar, except that fluorescently labeled Ub-Degron was used instead of Degron.

### Generation of the stable complex mimicking the transition state of the initiation step

#### 1. Preparation of Ub(1-75) hydrazide

Ub(1-75) hydrazide (Ub75-NHNH_2_) were generated using previously reported protein hydrazinolysis method ^42^. In brief, Ub(G76C) of which Gly76 at the C-terminus was mutated to Cys could undergo N-S acyl transfer so that hydrazine could be used as a suitable nucleophile, leading to a reliable C-terminal hydrazinolysis. 20 mg/mL Ub(G76C), 5 mg/mL TCEP, 50 mg/mL NHNH_2_•HCl, and 100 mg/mL MesNa were mixed in 20 mM Tris, pH 6.5, and stirred at 60 rpm and 50°C for 24 hours. The final products were purified and analyzed by RP-HPLC.

#### 2. Preparation of Ub(G76C)- ^K17^Degron

The reaction scheme was shown in Extended Data Fig. 3a. In brief, Ub75-NHNH_2_ peptide (1 μmol, 1 equiv) was dissolved in 1mL ligation buffer (6 M guanidinium chloride, 100 mM NaH_2_PO_4_, pH 2.3) precooled to −15 °C. Then 10 μL 1 M NaNO_2_ (10 μmol, 10 equiv) was added, and the reaction was stirred for 30 min at −15 °C to fully convert the hydrazide to the acyl azide. Next, sodium 2-mercaptoethanesulfonate (MesNa, 100 μmol, 100 equiv) was added, and the pH was adjusted to 5.0 for overnight reaction. The product, Ub75-MesNa, was further purified by RP-HPLC. Purified Ub1-75-MesNa (1 μmol, 1 equiv) and Cys-^K17^Degron peptide (1.1 μmol, 1.1 equiv) were mixed to the ligation buffer (6 M guanidinium chloride, 100 mM NaH_2_PO_4_, 5mg/mL TCEP, pH 7.4). Next, 4-mercaptophenylacetic acid (MPAA, 50 μmol, 50 equiv) was added, and the pH was adjusted to 6.4 for overnight reaction. The final product, Ub(G76C)- ^K17^Degron, was purified and analyzed by RP-HPLC.

#### 3. Preparation of Ubc2-Ub(G76C)- ^K17^Degron through disulfide ligation

Lyophilized dry power of Ub(G76C)- ^K17^Degron (1.3 mg) was dissolved in 100 μL 6 Mguanidinium chloride, 50 mM HEPES, pH 7.5 to a final concentration of 1mM. 2 μL of 100 mM 5,5′-dithiobis-(2-nitrobenzoic acid) (Sigma Aldrich, dissolved in 50 mM NaH_2_PO_4_, pH 7.5) was immediately added and fully mixed by pipetting before incubating at room temperature for 20 min. The solution was then diluted to the refolding buffer containing 50 mM HEPES pH7.5, 150mM NaCl. Excess reactants were removed by a Superdex peptide size-exclusion column (GE Healthcare) equilibrated in the refolding buffer. Finally, the product and 0.9 equiv Ubc2 (pre-dialyzed into the refolding buffer) were mixed and incubated at room temperature for 30 min. The final product was purified and analyzed by RP-HPLC.

### Generation of the stable complex mimicking the transition state of the elongation step

#### 1. Preparation of Molecule 2

As shown in Fig. 2b, **Molecule 2** was prepared using Cys-aminoethylation reaction^36^. Specifically, 1 μmol lyophilized powder of Ubc2 was incubated in aqueous alkylation buffer (6 M guanidinium chloride, 0.1 M HEPES pH 8.5, 5 mg/mL TCEP) with 40 mM **Molecule 1** (2-((2- chloroethyl)amino)ethane-1-(S-acetaminomethyl)thiol) at 37 °C for 14-16 h. The product, **Molecule 2**, was further purified by semi-preparative HPLC and lyophilized.

#### 2. Preparation of Molecule 3

Lyophilized **Molecule 2** was dissolved in reaction buffer containing 6 M guanidinium chloride,

0.1 M NaH_2_PO_4_ buffer, pH 7.4 at a final concentration of 1 mM. Then PdCl_2_ (15 equiv, pre-dissolved in the reaction buffer) was added and the mixture was incubated at 37°C for 1 h to remove the Acm group. Purified Ub(1-75)-MesNa (1 μmol, 1 equiv) and Molecule 2 (1.1 μmol, 1.1 equiv) were then mixed in the ligation buffer (6 M guanidinium chloride, 100 mM NaH_2_PO_4_, 5mg/ml TCEP, pH 7.4). Next, 4-mercaptophenylacetic acid (MPAA, 50 μmol, 50 equiv) was added, and the pH was adjusted to 6.4 to initiate native chemical ligation. The product was purified and analyzed by RP-HPLC.

#### 3. Preparation of Ub^K48C^-^K17^Degron

Different from the preparation strategy for Ub-Degron, we mutated the Lys48 to Cys, which enabled native chemical ligation. Furthermore, the thiol group on Cys48 was retained for disulfide ligation. Specifically, Ub(1-47)NHNH_2_ peptide (1 μmol, 1 equiv) was dissolved in 1mL ligation buffer (6 M guanidinium chloride, 100 mM NaH_2_PO_4_, pH 2.3) precooled to −15 °C. Then, 10 μL 1 M NaNO_2_ (10 μmol, 10 equiv) was added, and the reaction was stirred for 30 min at - 15 °C to fully convert the hydrazide to acyl azide. Next, Ub(48-76)^K48C^-^K17^Degron peptide (1.1 μmol, 1.1 equiv) was added to the ligation buffer, followed by 4-mercaptophenylacetic acid (MPAA, 50 μmol, 50 equiv). The pH was adjusted to 6.4 to initiate the ligation. The product, Ub^K48C^-^K17^Degron, was purified and analyzed by RP-HPLC.

#### 4. Preparation of Ubc2-Ub-Ub^K48C^-^K17^Degron using disulfide ligation

Lyophilized dry power of **Molecular 3** was dissolved in 500ul 6 M guanidinium chloride, 50mM HEPES, 1mM TCEP, pH7.5, and refolded through gradient dialysis against refolding buffer (50mM HEPES, 1mM TCEP, pH7.5) containing 6M, 2M, 1M to 0M guanidinium chloride. 1.3mg Lyophilized dry power of **Ub^K48C^-^K17^Degron** was dissolved in 100 μl 6 M guanidinium chloride, 50mM HEPES, pH7.5 (the final concentration was 1mM), and then 2μl 100mM 5,5′-dithiobis-(2-nitrobenzoic acid) (Sigma Aldrich, dissolved in 50 mM NaH_2_PO_4_ pH 7.5) was added and fully mixed by pipetting before incubating at room temperature for 20 min. The solution was then diluted to the refolding buffer, and the excess small molecular was removed by Superdex peptide size-exclusion column (GE Healthcare) equilibrated in refolding buffer. Finally, the pooled product and 1.1eq refolded **Molecular 3** were mixed and incubated at room temperature for 30 min. The final product, **Ubc2-Ub-Ub^K48C^-^K17^Degron**, was purified and analyzed by RP-HPLC.

### Specimen preparation for single-particle cryo-EM

Ubr1 (0.4 mg/mL) was mixed with 1.5-fold excess (molar ratio) initiation or elongation intermediate-mimics and incubated on ice for 30 min. A final concentration of 0.01% fluorinated octyl maltoside (FOM) was immediately added to the sample before grid freezing using a Vitrobot mark IV (Thermo Fisher) operating at 8 °C and 100% humidity. A volume of 3.5 μL sample was applied to a glow-discharged Quantifoil Cu 1.2/1.3 grid, and blotted for 1 second using standard Vitrobot filter paper (Ted Pella, 47000-100) before plunge freezing in liquid ethane.

### Data collection for single-particle cryo-EM

Optimized frozen grids were sent to Advanced Electron Microscopy Facility at the University of Chicago or National Cryo-Electron Microscopy Facility at National Cancer Institute for data collection. All datasets were acquired as movie stacks with a Titan Krios electron microscope (Thermo Scientific) operating at 300 kV, equipped with a Gatan K3 direct detection camera. A single stack consists of 40 frames with a total exposure around 50 electrons/Å**^2^**. The defocus range was set at −1.0 to −2.5 μm. See **Table S1** for the details.

### Image processing

Movie stacks were subjected to motion correction using MotionCor2 ^44^. CTF parameters for each micrograph were determined by CTFFIND4 ^45^. The following particle picking, two- and three-dimensional classifications, and three-dimensional refinement were performed in RELION- 3 ^23^. About 2,000 particles were manually picked to generate 2D class averages. The class averages were then used as templates for the following automatic particle picking. False-positive particles or particles classified in poorly defined classes were discarded after 2D classification. Initial 3D classification was performed on a binned dataset using the initial model obtained in RELION. The detailed data processing flows are shown in Extended Data Figs. 3 and 5. Data processing statistics are summarized in **Table S1**. Reported resolutions are based on Fourier shell correlation (FSC) using the FSC=0.143 criterion. Local resolution was determined using the implementation in RELION.

### Model Building, Refinement, and Validation

Ubr1 is a single-subunit E3 with more than 1,900 amino acids. Only was the structure of Ubr-Box1 domain determined previously ^24^. To overcome the difficulty during model building, we used artificial intelligence (AI) based de novo modelling tool DeepTracer program ^25^ to build a starting model of the entire complex from scratch. Specifically, the sharpened map of the initiation complex and a FASTA file containing sequences of Ubr1, Ubc2, and Ub were input to the online server of DeepTracer (https://deeptracer.uw.edu/home). The program finished within 10 minutes and output a complete model with Ubc2, and Ub correctly positioned. About 80% of Ubr1 was also correctly built into the cryo-EM map, with some errors in the poorly resolved regions and the zinc-binding sites. The starting model was first refined in real space using PHENIX ^27^, and then manually fixed, adjusted and refined using COOT ^26^. At the end, about 1800 residues of Ubr1 were built. The registration of the main chain was carefully checked and fixed based on bulky residues. The entire procedure was greatly simplified and accelerated with the starting model from DeepTracer. The final model was refined again in real space using PHENIX ^27^. The statistics of model refinement and geometry is shown in **Table S2**. Molecular graphics and analyses were performed with UCSF ChimeraX^46^ and ‘Protein interfaces, surfaces and assemblies’ service PISA at the European Bioinformatics Institute. (http://www.ebi.ac.uk/pdbe/prot_int/pistart.html) ^47^.

### Ubc2∼Ub thioester formation in the presence of U2BR peptide

First, 0.5 μM Uba1, 5 μM Ubc2, and 7.5 μM fluorescently labeled Ub was mixed in the reaction buffer containing 50 mM HEPES pH 8.0, and 150 mM NaCl. U2BR peptide was prepared as a twofold dilution series from the stock and added to the reaction mixture. Prepared ATP·Mg^2+^ mixture (50 mM MgCl_2_, 50 mM ATP, pH 8.0) was added to initiate the Ubc2∼Ub thioester formation. Reactions were quenched after 1 minute at 30 °C using 2 × SDS-sample buffer (pH < 3), followed by SDS-PAGE. The gels were imaged on ChemiDoc MP Imaging System, and substrate and product bands were quantified respectively as a percentage of the total signal for each time point using ImageLab (Bio-Rad). Ratio of ubiquitylated products relative to the total signal was plotted against the concentrations of Ubc2 and fit to the inhibitor vs. response model (three parameters) in GraphPad Prism 8.

### Isothermal titration calorimetry (ITC) analysis

All reported isothermal titration calorimetry data were collected using a MicroCal ITC 200 instrument in the center of biomedical analysis, Tsinghua University. Ubc2 and all U2BR peptides were buffer exchanged to 50 mM HEPES pH 7.5, and 150 mM NaCl before the experiment. For the experiments, 20 μM Ubc2 solution in the sample cell was titrated with 400 μM U2BR peptide solution through 19 injections (2.0 μL each) at 25 °C and 750 rpm stirring speed. Data fitting and analysis was performed using Origin 7 SR4 (OriginLab).

